# OpenMonkeyChallenge: Dataset and Benchmark Challenges for Pose Tracking of Non-human Primates

**DOI:** 10.1101/2021.09.08.459549

**Authors:** Yuan Yao, Abhiraj Mohan, Eliza Bliss-Moreau, Kristine Coleman, Sienna M. Freeman, Christopher J. Machado, Jessica Raper, Jan Zimmermann, Benjamin Y. Hayden, Hyun Soo Park

## Abstract

The ability to automatically track non-human primates as they move through the world is important for several subfields in biology and biomedicine. Inspired by the recent success of computer vision models enabled by benchmark challenges (e.g., object detection), we propose a new benchmark challenge called OpenMonkeyChallenge that facilitates collective community efforts through an annual competition to build generalizable non-human primate pose tracking models. To host the benchmark challenge, we provide a new public dataset consisting of 111,529 annotated (17 body landmarks) photographs of non-human primates in naturalistic contexts obtained from various sources including the internet, three National Primate Research Centers, and the Minnesota Zoo. Such annotated datasets will be used for the training and testing datasets to develop generalizable models with standardized evaluation metrics. We demonstrate the effectiveness of our dataset quantitatively by comparing it with existing datasets based on seven state-of-the-art pose tracking models.

## Introduction

Recent years have seen great advances in systems that can automatically track major landmarks in moving animals without fiducial markers, that is, *pose* (1–5). Such tracking systems have greatly benefited research in fields that study the tracked species (e.g., rodents, flies, and fishes). However, the ability to track non-human primates has lagged, rendering the primate order a major outstanding problem in the field (6). At the same time, non-human primates remain of great interest in biomedicine and related fields, including in neuroscience and psychology, as well as in anthropology, epidemiology, and ecology. Automated tracking can also benefit animal welfare programs, veterinary medical practice and, indeed, conservation projects.

Non-human primates (NHPs) are particularly challenging to track due to their homogeneous body texture and exponentially large pose configurations (6). Two major innovations are needed to solve the pose tracking problem in NHPs. (1) Algorithmic innovation: tracking models are expected to learn a generalizable visual representation that encodes the complex relationship between the visual appearance and spatial landmarks, which allows detecting poses in images with diverse primate identities, species, scenes, backgrounds, and poses in the wild environment. Existing deep learning models including convolutional pose machine (7), stacked hour-glass model (8), DeeperCut (9), and AlphaPose (10) incorporate a flexible representation with a large capacity, which have shown strong generalization on human subjects. However, these models are not applicable to the image samples of NHPs from the out-of-training-distribution due to their characteristics (homogeneous appearance and complex pose). (2) Data innovation: the tracking models learn the visual representation from a large annotated dataset that specifies the locations of landmarks. Existing publicly available datasets including OpenMonkeyPose (200K multiview macaque images in a specialized laboratory environment) (6) and MacaquePose (13K in-the-wild macaque images) (11) are important resources for the development of tracking algorithms, and as such, extend the boundary of pose tracking performance of NHPs. However, due to limited data diversity (appearance, pose, viewpoint, environment, and species), existing datasets are currently insufficient for learning generalizable tracking models.

Here we describe a novel dataset consisting of 111,529 images of NHPs in natural contexts with 17 landmark annotations. These datasets are obtained from various sources including the Internet, three National Primate Research Centers, and the Minnesota Zoo. Our motivation for developing this dataset includes inspiration from the recent success of computer vision models for human pose estimation (12), object detection (13), and visual question answering (14), enabled by standard benchmark challenges. For instance, the COCO benchmark challenges on object detection, segmentation, and localization have facilitated collective community effort through an annual competition, which in turn has been a driving force to advance computer vision models (13). In these domains, such datasets have served as a common comparison for friendly competitions, as a goal for experimentation, and as a benchmark to evaluate innovations. At the same time, such datasets tend to be difficult and expensive to generate, so sharing them makes economic sense for the field. Making them public greatly lowers the barriers to entry for new teams with innovative ideas.

With our dataset, we present a new benchmark challenge called *OpenMonkeyChallenge* for NHP pose tracking (http://openmonkeychallenge.com). It is an open and ongoing competition where the performance of each model is measured by the standard evaluation metrics. We leverage our unprecedentedly large annotated dataset, which includes diverse poses, species, appearances, and scenes as shown in Fig. 1. We split the dataset into the training and testing datasets where the testing dataset is used to evaluate the performance of competing models. We demonstrate that our dataset addresses the limitation on data diversity in the existing datasets. Specifically, we show the effectiveness of our dataset quantitatively by comparing it with existing datasets (e.g., OpenMonkeyPose and MacaquePose) based on state-of-the-art pose tracking models.

**Fig. 1.**
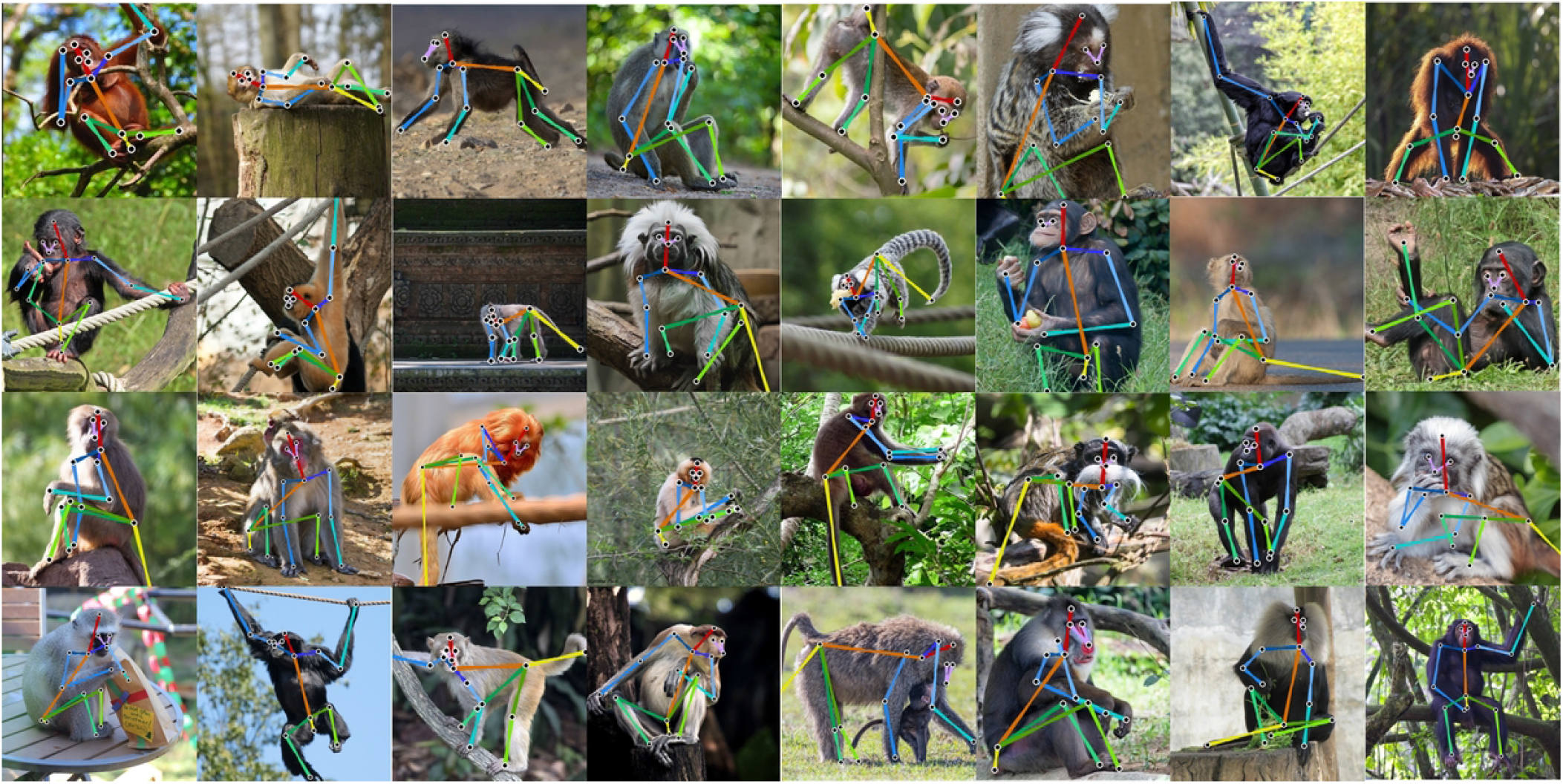
We present an OpenMonkeyChallenge using 111,529 annotated images of non-human primates (26 species), obtained from the internet, three National Primate Research Centers, and the Minnesota Zoo. 17 landmarks are manually annotated for each image. OpenMonkeyChallnege aims to extend the boundary of pose tracking for non-human primates across multiple species through an annual competition to build generalizable pose tracking models.

## Results

### OpenMonkeyChallenge and Benchmark Dataset

We collected 111,529 images of 26 species of primates (6 New World monkeys, 14 Old World monkeys, and 6 apes), including Japanese macaques, chimpanzees, and gorillas from (1) the internet images and videos, such as Flickr and YouTube, (2) photographs of multiple species of primates from three National Primate Research Centers, and (3) multiview videos of 27 Japanese macaques in the Minnesota Zoo (Fig. 2(b) and (d)). For each photograph, for example, in Fig. 1, we cropped the region of interest such that each cropped image contains at least one primate. We ensured that all cropped images have a higher resolution than 500×500 pixels.

**Fig. 2.**
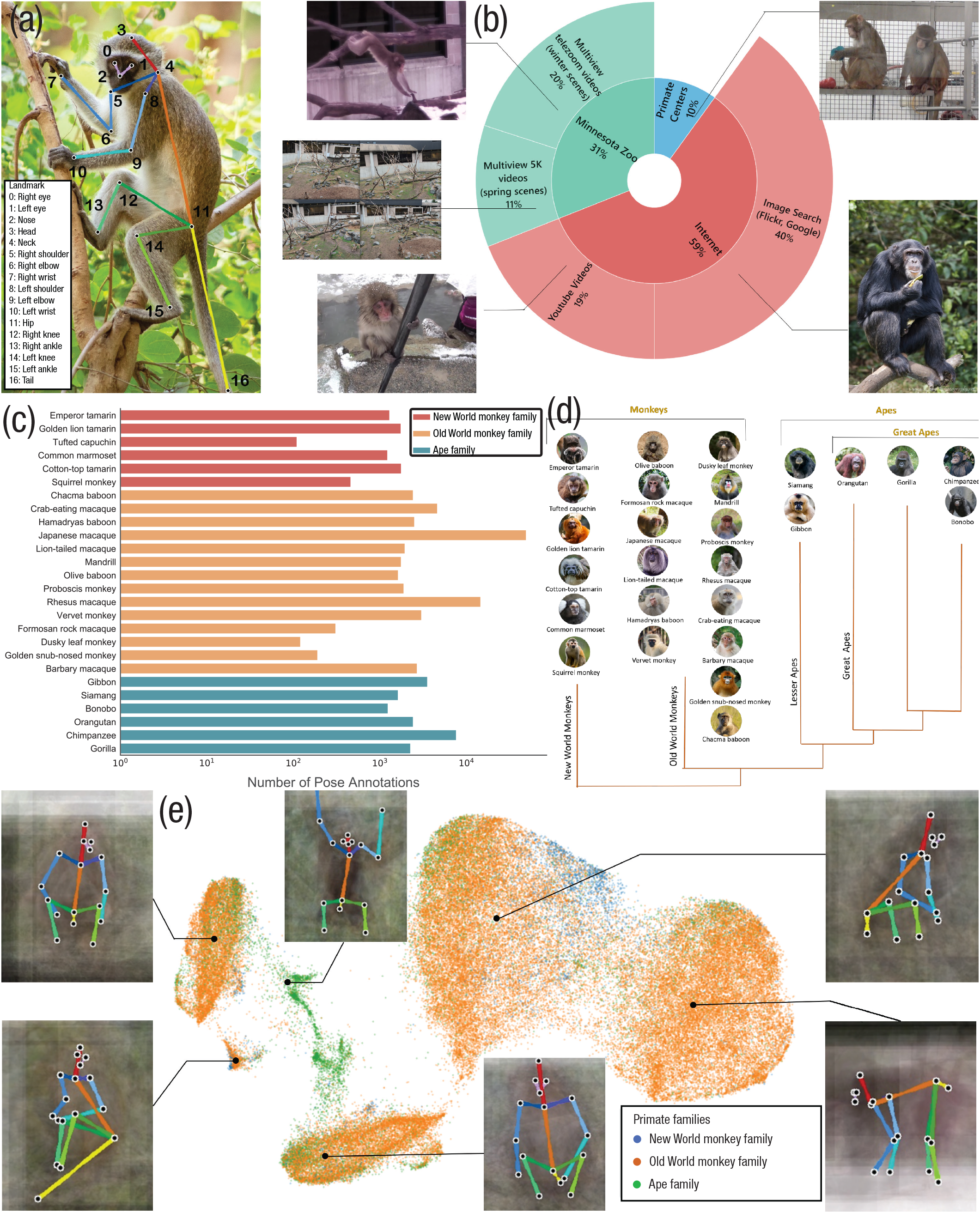
(a) We annotated the 17 landmarks that describe the pose of the primate in an image. (b) We collect image data from diverse sources: Internet image searches and YouTube videos, professional photographs from three National Primate Research Centers, and multiview videos from the Minnesota Zoo. The original images are cropped to include at least one primate and ensured to have higher than 500×500 resolution. (c) Our dataset is composed of 26 species of monkeys and apes, and more than 100 images are annotated for each species. We split the data into training, validation, and testing datasets, approximately 6:2:2 ratio, respectively. (d) Primate taxonomy. Our dataset includes diverse species of monkeys and apes. (e) We visualize a distribution of poses of the OpenMonkeyChallenge dataset using UMAP for dimension reduction. For each cluster, we show an average image overlaid with the median pose to illustrate its visual pattern.

We identify the region of interest (i.e., bounding box detection) by bootstrapping with a weak monkey detector (15) followed up by manual refinement and use a commercial annotation service (Hive AI) to manually annotate the 17 landmarks (See Method section.). The 17 landmarks together comprise a pose. Our landmarks include Nose, Left eye, Right eye, Head, Neck, Left shoulder, Left elbow, Left wrist, Right shoulder, Right elbow, Right wrist, Hip, Left knee, Left ankle, Right knee, Right ankle, and Tail. Each data instance is made of a triplet, image, species, pose as shown in Fig. 2(a).

We split the benchmark dataset into training (66,917 images, 60%), validation (22,306 images, 20%), and testing (22,306 images, 20%) datasets. We minimize visually similar image instances across splits by categorizing them using the time of capture, video and camera identification numbers, and photographers. Fig. 2(c) illustrates the data distribution across species, and each species includes more than 100 annotated images.

The annotations for the training and validation datasets are publicly available while that for the testing dataset is hidden. We have established the evaluation server to automatically evaluate the performance of the competing models on the testing dataset and maintain the leader. Specifically, the species landmark detection result on training/validation/testing datasets is uploaded to the evaluation server in a pre-defined file format, and the evaluation result is generated by the server. Users are asked to post their results in the leaderboard that sorts the performance based on three standard keypoint metrics: mean per joint position error (MPJPE), probability of correct keypoint (PCK) metric at error tolerance, and average precision (AP) based on object keypoint similarity (OKS).

Mean per joint position error (MPJPE) (16) measures normalized error between the detection and ground truth for each landmark (the smaller, the better):

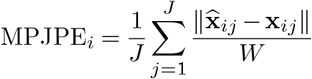

where MPJPE_*i*_ is the MPJPE for the *i*^th^ landmark, *J* is the number of image instances, 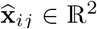 is the *i*^th^ predicted landmark in the *j*^th^ image x_*ij*_ ∈ ℝ^2^ is its ground truth location, and *W* is the width of the bounding box. Note that MPJPE measures the normalized error relative to the bounding box size *W*, e.g., 0.1 MPJPE for 500 × 500 bounding box corresponds to 50 pixel error.

Probability of correct keypoint (PCK) (17) is defined by the detection accuracy given error tolerance (the bigger, the better):

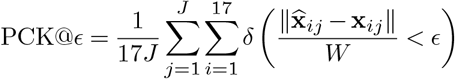

where *δ*(·) is an indicator function that outputs 1 if the statement is true and zero otherwise. *ϵ* is the spatial tolerance for correct detection. Note that PCK measures the detection accuracy given the normalized tolerance with respect to the bounding box width, e.g., PCK@0.2 with 200 pixel bounding box size refers to the detection accuracy where the detection with the error smaller than 40 pixels is considered as a correct detection.

Average precision (AP) measures detection precision (the bigger, the better):

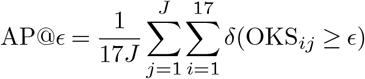

where OKS measures keypoint similarity (13):

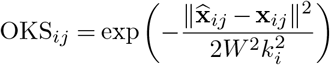

where OKS_*ij*_ is the keypoint similarity of the *j*^th^ image of the *i*^th^ landmark. *k*_*i*_ is the *i*^th^ landmark relative tolerance. Unlike PCK, OKS measures per landmark accuracy by taking into account per landmark variance *k*_*i*_ (visual ambiguity of landmarks), e.g., eye is visually less ambiguous than hip. We define *k*_*i*_ based on COCO keypoint challenge and augment the tail landmark such that *k*_*tail*_ = *k*_*wrist*_.

### Data Analysis

The OpenMonkeyChallenge dataset contains a diversity of species, poses, and appearances. We use Uniform Manifold Approximation and Projection (UMAP) (18) to reduce the high dimensional pose (ℝ^34^ for 17 landmarks) into two dimensions as shown in Fig. 2(e). To generate a spatially meaningful distribution, we normalize the pose coordinates. Specifically, the coordinates of each pose (17 landmarks) are normalized by centering the root landmark (hip joint), i.e., the landmark coordinate is relative with respect to the hip joint. These relative coordinates are normalized by the size of the bounding box to account for different sizes of images. Further, we align the orientation such that all poses have the same facing directions. This results in coherent clusters with poses.

The primates are classified into three types based on their families: New World monkeys, Old World monkeys, and apes. Poses are distributed across species, which are highly correlated with the semantically meaningful poses such as sitting, standing, and climbing. For each cluster, we visualize average images by aligning the poses. Overall, we find that the majority of data consists of sitting poses from a variety of views.

The clustering results also highlight the difference in locomotion patterns locomotion among primate families. For example, Old World monkeys (orange) heavily outnumber the other two families and dominate most of the clusters, and a few clusters of which the average pose is vertical climbing are by large composed of the apes (green). Other actions, such as sitting, walking, and standing, are common in all the primate families.

### Cross-dataset Evaluation

To evaluate the generalizability of our dataset, we conduct a cross-dataset evaluation with OpenMonkeyPose (6) and MacaquePose (11). OpenMonkeyPose (6) consists of 195,228 annotated images simultaneously captured by 62 precisely arranged high-resolution video cameras. The dataset involves inanimate objects (barrels, ropes, feeding stations), two background colors (beige and chroma-key green), and four rhesus macaque subjects varying in size and age (5.5–12 kg). MacaquePose (11), a dataset with more than 13,083 images of macaque, is collected by searching for images with a ‘macaque’ tag in Google Open Images and captured in zoos and the Primate Research Institute of Kyoto University.

We split each dataset into training (60%), validation (20%), and testing (20%) sets. We train a convolutional pose machine (CPM) (7) using the training data from one of the datasets with spatial data augmentation (translation and rotation) until it starts to overfit based on the model performance on the validation data, and test that model on the testing data from each dataset. Fig. 3 summarizes the performance in MPJPE. The CPM model trained by the OpenMonkeyChallenge dataset achieves the lowest MPJPE on the OpenMonkeyChallenge and MacaquePose (11) test datasets, which indicates that the diversity and generalizability of our training dataset (outperforming MacaquePose own testing data). For the OpenMonkeyPose testing dataset, it achieves the second best close to the OpenMonkeyPose. This is mainly caused by the domain difference: the images of OpenMonkeyStudio were captured by a controlled lab environment that has a homogeneous background and monkey texture. For the same reason, this model has poor performance on the other two datasets due to its low generalizability.

**Fig. 3.**
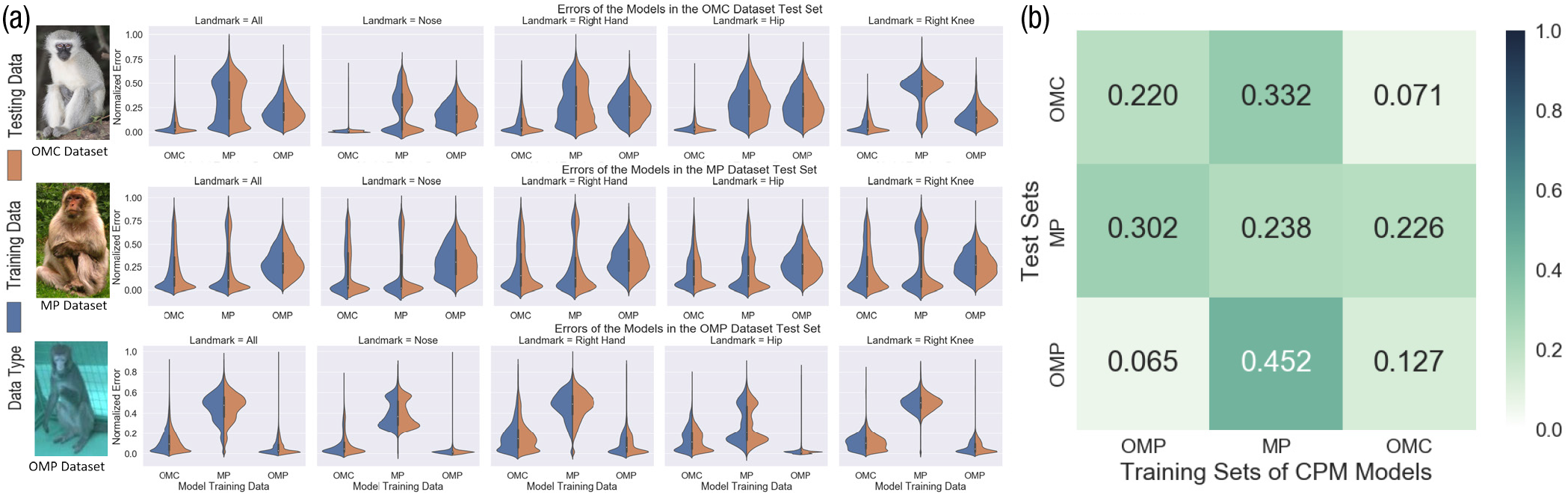
(a) Three detection models are trained on OpenMonkeyChallenge (OMC), MacaquePose (MP), and OpenMonkeyPose (OMP), respectively. In each box, we visualize three violin plots corresponding to the detection models. Each violin plot shows the normalized error histogram of landmarks on training (blue) and testing (brown) data (first row: OMC dataset; second row: MP dataset; third row: OMP dataset). The model trained on OMC (left violin plot in each box) is the most generalizable (inverted T shape histogram). (b) We summarize cross-dataset evaluation to show the generalizability using the normalized error in a confusion matrix, e.g., the second row of the third column shows the normalized error of the MP testing data for the model trained on OMC training dataset. The model trained on OMC dataset shows the smallest error or comparable to the model that is testing on its own training data.

### Comparison with Human Pose Estimation

The distal goal of our benchmark challenge is to achieve a performance comparable to human pose estimation. For instance, a state-of-the-art human pose detector (CPM) trained on the COCO-keypoint dataset (13) produces 0.061 MPJPE or 0.849 PCK@0.2 (upper bound performance). Without a non-trivial modification, a CPM trained on our dataset achieves 0.074 MPJPE or 0.761 PCK@0.2 as reported in Fig. 4(a). In other words, there exists a considerable performance gap between human and primate pose estimation. Further, we show the human detection model on out dataset, which achieves 0.197 MPJPE or 0.265 PCK@0.2 for reference (lower bound performance). We propose that the major benefits associated with human pose estimation is the progress in developing, efficient and generalizable models with self-supervised methods (19–24). We anticipate that a similar algorithmic innovation will close the gap.

**Fig. 4.**
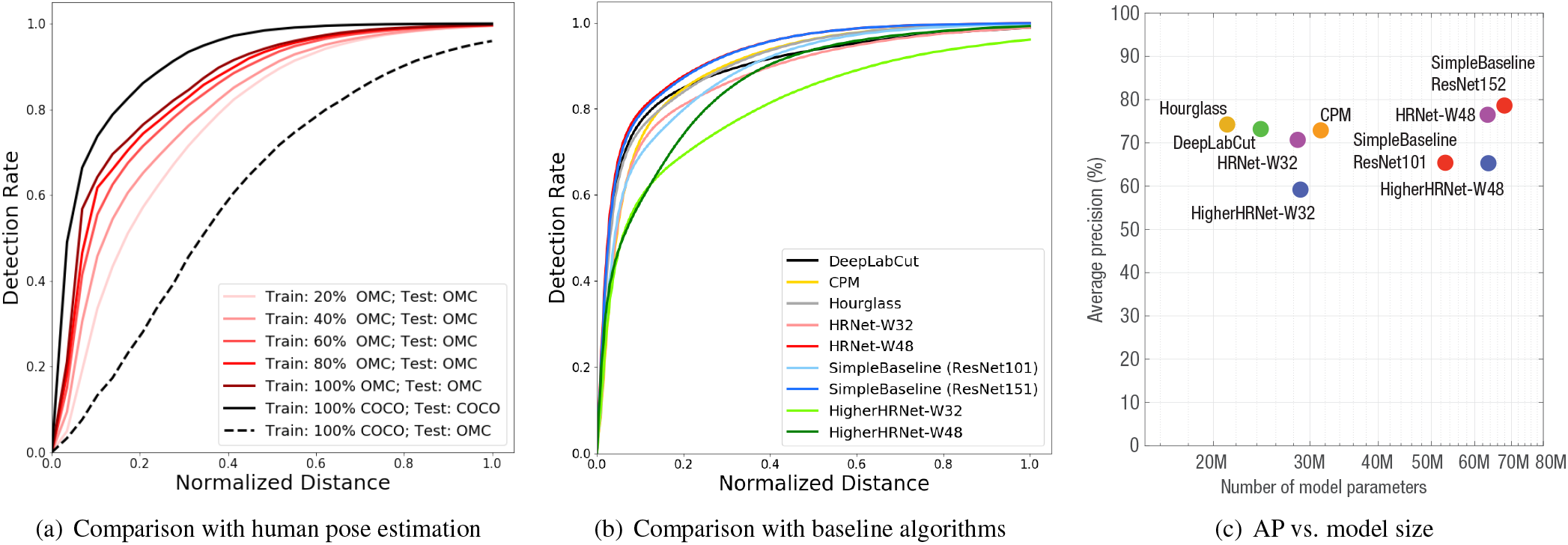
(a) We use PCK to measure keypoint detection performance. The black solid line shows the performance of the human landmark detector (train and test on COCO) that forms the upper bound of the primate landmark detector. The black dotted line shows the testing performance of the human landmark detector (trained on COCO) on OMC data without retraining, which forms the lower bound. OMC dataset allows us to train a primate specific model that shows significant performance improvement from the lower bound. Yet, there still exists a large gap between the human and primate landmark detectors. We also visualize the performance improvement as increasing the number of OMC training data. (b) Six state-of-the art pose estimation models are trained with OMC datasets. These are PCK curves in the test set from these models. (c) We show the average precision (AP) of state-of-the-art models as a function of the number of model parameters. If the data size is large enough, a larger model is likely to learn complex visual patterns.

**Fig. 5.**
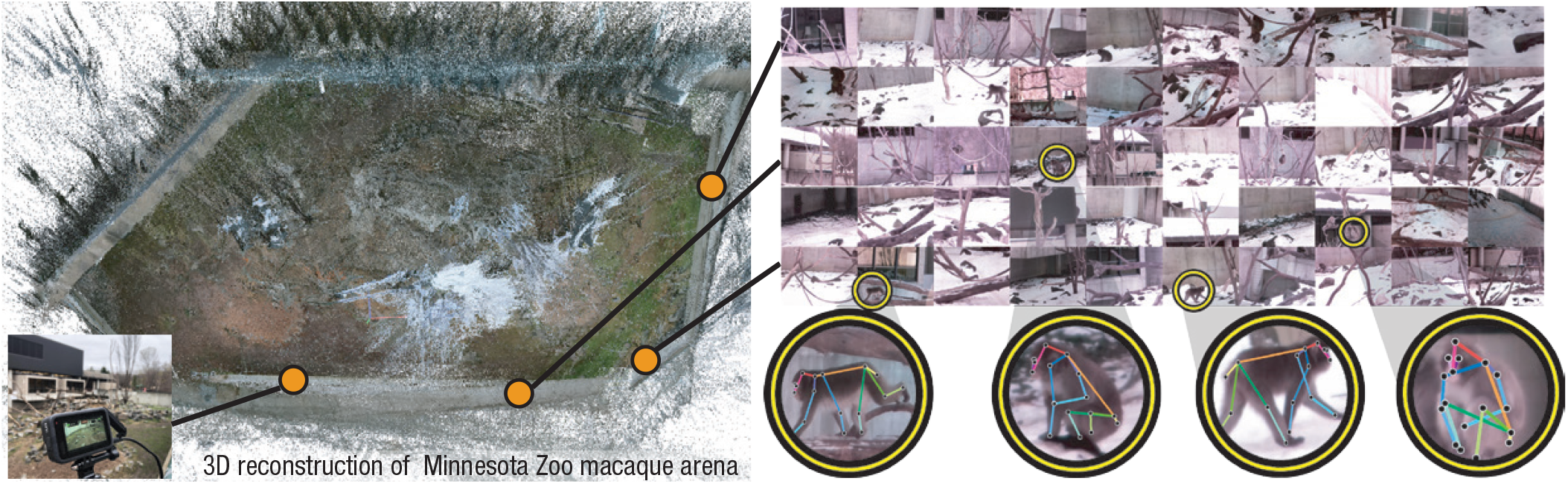
We design a graphic user interface to refine bounding boxes. Given an image and bounding box proposals (green boxes) from a weak detector, the annotators are asked to remove false positives and redundant poses from previous frames (green bounding boxes with red cross) and to add false negatives (red bounding boxes).

Further, we conduct an ablation study to evaluate the impact of large data, i.e., how the amount of training data affects the landmark detection accuracy on the testing dataset. Given the training data, we incrementally reduce the amount of the training images used for model training by 20% at each time and measure the model performance using PCK metric. Fig. 4(a) shows the impact of the data increments, i.e., the model trained on 100% training data achieves the highest PCK result, outperforming the model with 20% of training data by 15% at PCK@0.2.

### State-of-the-art Detection Model Performance Evaluation

We conduct a comparative study on the performance of the state-of-the-art pose detection models using the OpenMonkeyChallenge dataset. We train nine pose estimation models until it starts to overfit based on the performance on the validation data. These models can be categorized into the top-down methods and the bottom-up methods. The top-down models (DeepLabCut with ResNet (25), CPM (7), Hourglass (8), HRNet-W32 (26), HRNet-W48, SimpleBaseline with ResNet101 (27), and SimpleBaseline with ResNet152) detect the keypoints of a single primate given the bounding box. In contrast, the bottom-up models (HigherHRNet-W32 (28) and HigherHRNet-W48) localize the landmarks without a bounding box and group them to form poses, specialized for multi-primate detection. For all models, we use their own pretrained model and training procedural protocol, i.e., the DeepLabCut model is pre-trained on ImageNet. The top-down models, in general, show stronger performance because of resolution while it shows weaker performance when multiple primates are present. Table 1 summarizes the normalized MPJPE of each landmark in the testing dataset predicted by six models across models. Table 2 reports the PCK@0.2 of each landmark in the testing dataset, and Fig. 4(b) shows the PCK curve of each model. In short, there is no clear winner. All models use a variant of high capacity convolutional neural networks that can effectively memorize and generalize the training data through fully supervised learning. SimpleBaseline (27) slightly out-performs other models (the lowest MPJPE and the highest PCK@0.2). Fig. 4(c) shows AP comparison as a function of the model parameters. In general, when the number of data is sufficiently large, larger and deeper models outperforms small and shallow models because more complex visual patterns can be learned.

**Table 1.**
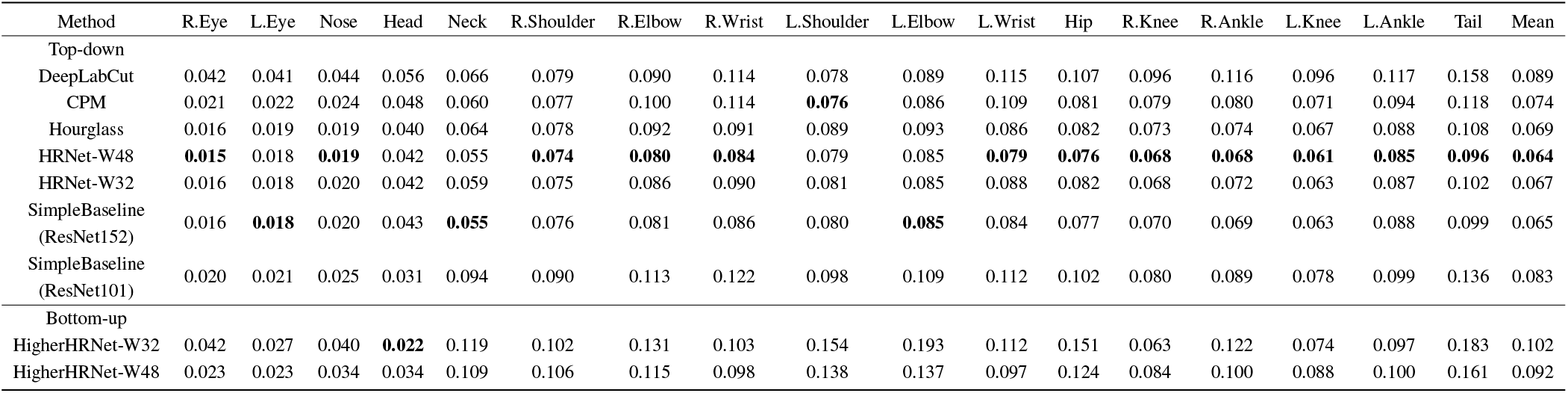
Model comparison with MPJPE metric of each landmark with top-down and bottom-up methods on the OpenMonkeyChallenge test set.

**Table 2.** Model comparison with PCK@0.2 metric of each landmark with top-down and bottom-up methods on the OpenMonkeyChallenge test set.

| Method | R.Eye | L.Eye | Nose | Head | Neck | R.Shoulder | R.Elbow | R.Wrist | L.Shoulder | L.Elbow | L.Wrist | Hip | R.Knee | R.Ankle | L.Knee | L.Ankle | Tail | Mean |
| --- | --- | --- | --- | --- | --- | --- | --- | --- | --- | --- | --- | --- | --- | --- | --- | --- | --- | --- |
| Top-down |  |  |  |  |  |  |  |  |  |  |  |  |  |  |  |  |  |  |
| DeepLabCut | 0.937 | 0.938 | 0.936 | 0.926 | 0.922 | **0.906** | **0.874** | 0.814 | **0.908** | **0.875** | 0.810 | 0.855 | 0.868 | 0.819 | 0.862 | 0.816 | 0.747 | 0.871 |
| CPM | 0.995 | 0.995 | 0.994 | 0.960 | **0.945** | 0.887 | 0.825 | 0.785 | 0.886 | 0.869 | 0.814 | 0.892 | 0.893 | 0.902 | **0.924** | 0.854 | 0.809 | 0.896 |
| Hourglass | 0.997 | 0.996 | 0.996 | 0.960 | 0.925 | 0.864 | 0.823 | 0.825 | 0.839 | 0.836 | 0.846 | 0.869 | 0.881 | 0.891 | 0.910 | 0.853 | 0.814 | 0.890 |
| HRNet-W48 | 0.997 | 0.996 | 0.996 | 0.951 | 0.940 | 0.876 | 0.858 | **0.846** | 0.867 | 0.856 | **0.864** | 0.885 | 0.893 | 0.906 | 0.923 | **0.863** | **0.842** | **0.903** |
| HRNet-W32 | 0.997 | **0.997** | 0.996 | 0.958 | 0.934 | 0.874 | 0.844 | 0.828 | 0.860 | 0.857 | 0.844 | 0.867 | 0.898 | 0.897 | 0.921 | 0.854 | 0.830 | 0.897 |
| SimpleBaseline (ResNet152) | **0.997** | 0.996 | **0.996** | 0.954 | 0.942 | 0.868 | 0.855 | 0.842 | 0.864 | 0.858 | 0.855 | 0.883 | **0.892** | **0.907** | 0.921 | 0.855 | 0.838 | 0.901 |
| SimpleBaseline (ResNet101) | 0.995 | 0.995 | 0.994 | **0.983** | 0.877 | 0.837 | 0.776 | 0.757 | 0.820 | 0.798 | 0.794 | 0.827 | 0.870 | 0.868 | 0.897 | 0.828 | 0.756 | 0.863 |
| Bottom-up |  |  |  |  |  |  |  |  |  |  |  |  |  |  |  |  |  |  |
| HigherHRNet-W32 | 0.962 | 0.979 | 0.981 | 0.978 | 0.856 | 0.798 | 0.724 | 0.782 | 0.661 | 0.620 | 0.804 | 0.710 | 0.899 | 0.804 | 0.897 | 0.820 | 0.633 | 0.818 |
| HigherHRNet-W48 | 0.986 | 0.986 | 0.985 | 0.965 | 0.860 | 0.796 | 0.777 | 0.820 | 0.721 | 0.730 | 0.831 | 0.779 | 0.863 | 0.834 | 0.874 | 0.827 | 0.715 | 0.844 |

**Table 3.** Model comparison with AP metric based on OKS of each landmark with top-down and bottom-up methods on the OpenMonkeyChallenge test set.

| Method | # Params | AP@0.5 | AP@0.6 | AP@0.7 | AP@0.8 | AP@0.9 | AP |
| --- | --- | --- | --- | --- | --- | --- | --- |
| Top-down |  |  |  |  |  |  |  |
| DeepLabCut | 24.4M | 92.3 | **89.7** | **83.9** | 74.1 | 52.6 | 73.2 |
| CPM | 31.4M | 91.8 | 86.1 | 78.9 | 69.6 | 54.2 | 72.9 |
| Hourglass | 21M | 91.3 | 85.7 | 80.8 | 74.7 | 63.8 | 74.5 |
| HRNet-W32 | 28.5M | 89.4 | 80.6 | 71.0 | 64.6 | 65.7 | 70.7 |
| HRNet-W48 | 63.6M | 90.2 | 85.7 | 80.8 | 74.7 | 63.8 | 76.5 |
| SimpleBaseline (ResNet152) | 68.0M | 89.5 | 84.9 | 81.2 | **76.9** | **67.8** | **78.5** |
| SimpleBaseline (ResNet101) | 53.0M | **97.2** | 82.6 | 65.9 | 46.9 | 31.4 | 65.3 |
| Bottom-up |  |  |  |  |  |  |  |
| HigherHRNet-W32 | 28.6M | 88 | 79.5 | 57.3 | 32.0 | 20.0 | 59.1 |
| HigherHRNet-W48 | 63.8M | 91.5 | 82.6 | 65.9 | 46.9 | 31.4 | 65.3 |

## Discussion

Here we present a new resource, a very large (111,529 images of 26 species) and fully annotated database of photographs of non-human primates. The primates come in a range of species and poses, and with a range of backgrounds. The primary goal of this resource is to serve as a training tool for scholars interested in developing computer vision approaches to identifying pose in the primate order. This resource can be found on our new website (http://openmonkeychallenge.com). The website also presents a new benchmark challenge for primate landmark detection. In parallel with our resource and the challenge, and as a baseline for modeling efforts, we provide some analyses of existing models. These analyses reveal that non-human primate detectors have substantially worse performance than human ones. We propose that our large dataset will be a critical tool in closing that performance gap.

We know of only two existing large datasets of annotated primate images, OpenMonkeyPose (6) and MacaquePose (11). OpenMonkeyPose, which our group developed, consists of nearly 200,000 annotated (13 landmarks) multiview (62 cameras) images of rhesus macaques in a specific carefully controlled laboratory environment. That dataset has a very different purpose than the present one—its chief virtue is its robust characterization of a single environment and species, and its multiview aspect for 3D motion capture. However, it is highly limited for the general purpose of pose identification because of its narrow number of backgrounds, species, individuals, and poses. The MacaquePose dataset, which consists of 13,000 images, is likewise limited to a single species and is also substantially smaller. Our analyses confirm that these datasets cannot be used to train robust models that can identify pose in general contexts nearly as well as this one can. These results, then, argue for the value of large variegated datasets like the one we present here. More generally, they demonstrate the critical importance of variety when training robust detection networks.

A key finding from our comparative study is that the state-of-the-art designs of convolutional neural networks (CNNs), including DeepLabCut, perform, by large, on a par with each other. These CNNs effectively learn a visual representation of primates from sufficiently large and diverse image data in a fully supervised manner where generalizable image features can be learned. This closes the gap between models. On the other hand, this finding implies that there is a fundamental limitation to the supervised learning paradigm. That is, our results indicate that the CNN models overfit to the training data; the distribution of the training data differs considerably from that of the testing data. As a consequence, the generalization is strictly bounded, which leaves a large performance gap between human and primate landmark detections. This requires employing the new semi- or unsupervised learning paradigm, which allows utilizing a potentially unlimited amount of unlabeled, or weakly labeled primate images, which can close the domain difference.

Through the OpenMonkeyChallenge, we aim to derive two major innovations to solve challenging computer vision problems. First, algorithmic innovation can lead to substantial performance gain by learning an efficient representation from a limited annotated data. Transfer learning, or domain adaptation, used in DeepLabCut is one of such kinds that leverage a pre-trained generic model learned from a large dataset (e.g., ImageNet). Such approaches have shown a remarkable generalization over frames within a target video while showing limited performance when applying to new videos with different viewpoints, poses, illumination, background, and identities. Second, data innovation can lead to great advances in generalization by large agnostic to algorithms and representations. For example, the field has witnessed such gains from the object detection community, e.g., from a few hundreds of images in Caltech-101 and Pascal VOC datasets to millions of images in ImageNet and COCO datasets (29). OpenMonkeyChallenge facilitates these two indispensable innovations for developing a generalizable primate detector through community effort.

This database and associated analyses are likely to be important for any fields in which behavioral tracking is important. We are especially sanguine about their potential to have positive benefits in neuroscience. Rhesus macaques are an important animal model in neuroscience, and other primate species are of growing importance as well. Their importance is unlikely to diminish, because of the unique anatomical homologies between members of the primate order, including many homologies between monkey and humans which rodents lack. While neuroscience does not, with a few notable exceptions, make use of primate tracking, the ability to track primates is of great interest in the field for several reasons. First, brains evolved hand in hand with our bodies and with the world in which they are embodied; there is a general sense that nothing in neuroscience makes sense except in light of behavior (30–33). Second, behavior represents a high-dimensional output that can give insight into inner cognitive states that are typically only viewed through a narrow aperture (such as reaction times or pupil size). As such, behavior promises to greatly expand the range of states that can be detected and linked to neural activity. Moreover, behavior can potentially reduce the need for neural measures, since it can serve as an alternative window into internal states. Finally, there is evidence that behavior drives neural activity to a major and heretofore unanticipated extent (34, 35). Measuring it can both allow us to regress out its effects and explain unexplained variance in firing, but more importantly, it can point us in new unanticipated directions. Indeed, it is quite possible that behavior is one of the major driving forces of neural activity, and so quantifying it can alter theories of brain activity.

## ACKNOWLEDGEMENTS

We thank Praneet Bala, Lin Huynh, Peeyush Samba, Justin Aronson, and Jen Holmberg for help on image acquisition. We thank the staff at the Minnesota Zoo for copious help, especially Tom Ness, Kathy Schlegel, Jamie Toste, Laurie Trechsel, and Kelli Gabrielson.

NSF IIS 2024581 to HSP, JZ, and BYH

NIH P51 OD011092 to ONPRC

NIH P51 OD011132 to YNPRC

R01-NS120182 to JR

K99-MH083883 to CJM

## Method

### Image Data Collection

We collected images from three sources: internet images and videos, photographs from National Primate Research Centers, and multiview videos from the Minnesota Zoo.

#### Internet images

Approximately 59% of our dataset were collected from the Internet through Image and video search engines. For instance, we used the Flickr API to scrape the list of image URLs and YouTube search engine to find relevant videos using species name keywords. We ensure visual diversity (shapes, poses, viewpoints, sizes, colors, and environments) and quality (image resolution, blurriness, lighting, and occlusion) of the scraped data via manual inspection. For the common species such as rhesus macaque, mandrill, and gorilla, image searches were sufficient. For the rarer species such as marmoset, we leveraged the video search features and extracted image frames from the videos. Not only does this approach allow us to obtain more images of the rarer species, but we also collected images that are less iconic than those from search engines. We hired two annotators for image and video searches. After image collections, we annotated the bounding boxes that contain the primate instances. For a subset of internet images, we do not own the copyright of the images. We specify the terms and conditions of use in the website.

#### Photographs from national primate centers

We made use of high quality images of primates photographed by staff at two National Primate Centers: Yerkes National Primate Research Center and the Oregon National Primate Research Center. The photographers were asked to capture primate images from diverse viewpoints and poses at high resolution (>2K pixel resolution) and often made use of a tele-zoom lens. 10,500 images are captured from the professional photographers across the primate centers. Further, we collected videos from California National Primate Research Center. Still images were extracted from a video library developed at the California National Primate Research Center (36, 37). Video footage of monkeys behaving was recorded at the center’s large 0.5 acre outdoor enclosures and from images of monkeys in the laboratory. Videos were edited to be 30 seconds in duration and included a range of behaviors, including aggression, grooming, feeding, resting, and affective displays. Still images were captured from the videos for use in this project.

#### Multiview videos from the Minnesota Zoo

We used video cameras to capture video images of a large troop (n=27 individuals) of snow monkeys (*Macaca fuscata*) at the Minnesota Zoo (Apple Valley, MN) for a long duration (1 week). Unlike the images taken by photographers who precisely control focal length and viewpoint to ensure high resolution images, these video cameras passively observe the scene. The monkeys inhabit a large arena that facilitates natural social interactions among them. It is a large open space (bigger than 600 m^2^), which leads to a new challenge as monkeys appear small in images (10-50 pixel size) if a wide field of view lens is used to cover the large area. We address this challenge by using a multi-camera system made of 20-30 cameras where each camera observes a small area (up to 5m×5m) using a narrow field of view (long or tele-zoom focal length). We identified the regions of the enclosure that frequently involve diverse activities (e.g., trails, ponds, and playgrounds) to maximize the monkey appearance in images. Because videos were multiview videos, we used a monkey bounding box detection algorithm to identify the monkeys and then refined these boxes manually. We collected the image data from two seasons (winter and spring) to maximize diversity of background visual appearance.

### Semi-automatic Annotation

Identifying images that contain primate instances from videos and annotating their landmarks are prohibitively labor intensive tasks. For instance, fewer than 2% of the frames in the videos from narrow field of view (FOV) cameras used in the zoo data contain primate instances. Watching every frame in videos to annotate bounding boxes for primate instances is time-consuming, e.g., one day zoo videos is equivalent to approximately 5,000 hours (∼6,000,000 images) of labor. Instead, we leverage an iterative bootstrapping approach to address the bounding box annotation task.

#### Bounding box proposal

We trained a weak primate detector that can predict the bounding box of a primate instance given an image. The bounding box (left-top corner coordinate, width, and height) of 3,000 internet images are manually annotated, and used to train a YOLOv3 model (15) that can recognize primate bounding boxes. We use a lower threshold for bounding box detection such that the false positives are slightly more common than the false negatives. This bounding box prediction automates identifying image frames that contain primate instances, so that a majority of image frames without primates can be pruned, which significantly reduces the required labor. Further, it provides bounding box candidates for each image.

#### Bounding box refinement

Given the bounding box proposals, we designed a graphic user interface to visualize and refine bounding boxes as shown in Fig. 6. The interface shows an image with bounding box candidates. The annotators are asked to find false positives and redundant poses from the previous frames (green bounding box with red cross). Further, they can add bounding boxes (red bounding boxes). Human helpers can perform this task in 5∼15 seconds per image. With this manual refinement, we ensure all cropped images include at least one primate. Once we have refined bounding box refinement, we incrementally increase the size of data to re-train the bounding box detection model to adapt to the target environments.

**Fig. 6.**
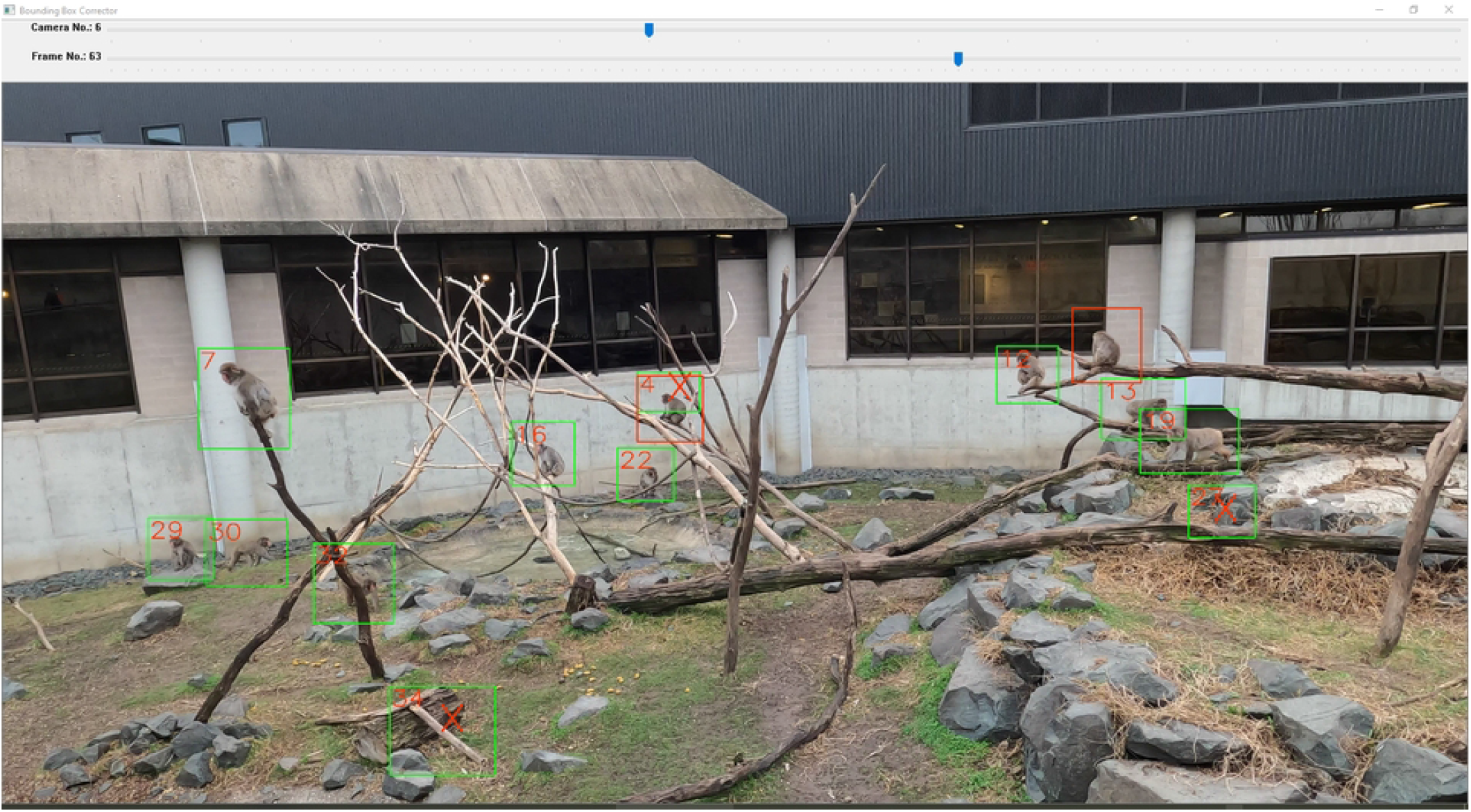
We design a graphic user interface to refine bounding boxes. Given an image and bounding box proposals (green boxes) from a weak detector, the annotators are asked to remove false positives and redundant poses from previous frames (green bounding boxes with red cross) and to add false negatives (red bounding boxes).

#### Landmark annotation

Given the bounding box annotations, we used a commercial annotation service (Hive AI) to annotate 17 landmarks from cropped images. When the landmarks are occluded, the annotators are instructed to specify the best guess location and to indicate visibility.

### Benchmark Evaluation Process

We created a website http://openmonkeychallenge.com/ that shares the dataset and benchmark challenges. The training/validation/testing datasets can be downloaded from the website. The annotations are available for the training and validation datasets. The testing results (landmark detection on the testing data) from the developed models can be submitted to the evaluation server in JSON file format:

~~~
   {“image_id” = int,
   “file_name” = str,
   “landmarks” = [x1,y1,…,x17,y17]}
~~~

where *x*_*i*_ and *y*_*i*_ are *x, y* coordinates of the *i*^*th*^ landmark. The evaluation server will return the performance on the testing data using MPJPE, PCK, and AP metrics. The evaluation results will be posted in the leaderboard that sorts the algorithms based on the performance. Optionally, the users can opt out. The website includes step-by-step description of the evaluation process, file format, and visualization code.

